# Study of growth characteristics of callus, suspension and root crops *Scutellaria baicalensis*

**DOI:** 10.1101/2021.06.06.447294

**Authors:** Anastasia Igorevna Dmitrieva, Anastasia Michailovna Fedorova, Violetta Mironovna Le

## Abstract

*Scutellaria baicalensis* is a popular traditional plant in Chinese medicine. Widely distributed biologically active substances of *Scutellaria baicalensis* are flavonoids (baicalin, baicalein, vogonin and vogonoside), which are responsible for the antitumor activity. Their antitumor effect is due to the absorption of oxidative radicals, the weakening of the activity of NF-kB (nuclear factor-kB), the suppression of the expression of the COX-2 gene and the regulation of the cell cycle. In addition, baicalein, baicalin, and vogonin showed strong antioxidant activity. The root culture in vitro of the medicinal plant Scutellaria baicalensis is characterized by intensive growth. The growth index at the end of the cultivation cycle was 40. The growth curve has a standard S-shape, with pronounced growth phases. The stationary phase was observed from 5-7 weeks of cultivation. The root culture growth index was 22.

## Introduction

*Scutellaria baicalensis* is a popular traditional plant in Chinese medicine. [1] Scientists report that the Baikal compounds contained in the skullcap extract have a wide range of antitumor activity both *in vitro* and *in vivo* (liver cancer, stomach cancer, lung cancer, breast cancer, prostate cancer, bladder cancer, brain cancer, squamous cell carcinoma, mucoepidermoid carcinoma, colorectal cancer, gallbladder cancer, oral cancer, leukemia, lymphoma and myeloma) [1].

Widely distributed BAS of skullcap are flavonoids (baicalin, baicalein, vogonin and vogonoside) [3], which are responsible for the antitumor activity of the plant. Their antitumor effect is due to the absorption of oxidative radicals, the weakening of the activity of NF-kB (nuclear factor-kB), the suppression of the expression of the COX-2 gene and the regulation of the cell cycle. In addition, baicalein, baicalin, and vogonin showed strong antioxidant activity [1].

It is known that baicalein is used in Asian medicine (in China and Japan) for the treatment of ischemic diseases [6]. It has also been proven that this compound has antioxidant activity. In the work of J. Y. Jeong, a study was conducted on the presence of the protective effect of baicalein on DNA damage and apoptosis, as a result of which, scientists proved that baicalein effectively inhibited H2O2-induced cytotoxicity and DNA damage by inhibiting the accumulation of reactive oxygen species (ROS) [2].

Vogonin-5,7-dihydroxy-8-methoxyflavone, a flavonoid-like chemical compound, is a flavone that has an antitumor effect, which consists in inhibiting the growth of cancer cells, by stimulating autophagic and apoptotic cell death [7].

Another valuable flavonoid is oroxylin A, which induces apoptosis in human colon cancer cells via the mitochondrial pathway [5], which has an anti-hepatic effect. In the study of H. Jin, it was proved that oroxylin A relieved alcoholic liver damage by inhibiting the aging of hepatocytes, in addition, it was proved that this compound also has anti-inflammatory, anti-cancer, antibacterial and antiviral effects [4].

### Objects and methods of research

#### Objects of research

– seeds of the Baikal skullcap (*Scutellaria baicalensis*), obtained and sprouted in the Botanical Garden of the Baltic Federal University named after I. Kant, Kaliningrad;
– callus, root and suspension cultures of Baikal skullcap (*Scutellaria baicalensis*);

For the obtained selected steadily growing callus cell cultures of medicinal plants of the Baikal skullcap (*Scutellaria baicalensis*), a preliminary analysis of the growth of the cultures was performed and the growth index was calculated.

At the first stage of studying the growth index of callus cultures, the callus cell cultures were characterized by studying the weight of raw biomass.

To determine the initial weight of the culture (the weight of the transplant), it was weighed before transplanting into a culture vessel (Petri dish) with a medium. After placing the graft on the medium, a second weighing was performed and the weight of the graft was determined as the difference between the second and first weighing. The weight of the grafts was equalized with an accuracy of ± 10 %. For callus cell cultures the weight of the graft was 0.1 g.

To determine the weight of the raw biomass of the culture during cultivation, the culture was removed from the culture vessel and weighed on an analytical balance.

Based on the results obtained, the increase in Pi biomass over a certain growing time (on the first day of cultivation) was determined by the formula 2.3.1:

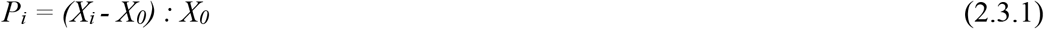

*Where:*

*X*_*i*_ - the weight of the crop on the i-th day of cultivation (standard: i = 7, 14, 21, 28 and 35 days of cultivation);

*X*_*0*_ - the initial weight of the culture (the weight of the transplant).

Next, we studied the growth characteristics of suspension cultures of medicinal plants of the Baikal skullcap (*Scutellaria baicalensis*).

To do this, we studied the physiological parameters of the obtained lines of primary relatively steadily growing suspension cell cultures - at 4-5 passages with standard periodic cultivation in flasks on a shaking machine under constant conditions. This method of cultivation is the most common for maintaining suspension cultures of plant cells in laboratory collections, and the physiological parameters of the strains obtained in the process of such cultivation are usually taken as control.

For the most stably growing suspension lines, the growth characteristics were comprehensively studied for all the parameters studied (dry weight, cell viability).

To determine the content of raw and dry biomass in a liter of medium, a fixed volume of suspension (not less than 15 ml, in three replications) was filtered through a paper filter using a Buchner funnel under vacuum. The biomass was dried to a constant weight in a current of air at a temperature of 30 ° C.

The viability of cell cultures was determined using the in vivo dye phenosafranin (0.1% solution), or 0.025 % Evans blue, by counting live (unpainted) and dead (colored) cultured units under a microscope.

Based on the results obtained, the growth index (I), the specific growth rate in the exponential phase (µ), the economic coefficient (Y), and the doubling time (τ) were calculated using the following formulas 2.3.2 and 2.3.3:

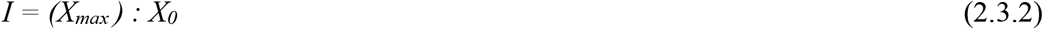

*Where:*

*X*_*max*_ - the maximum value of one of the growth criteria (in this work – the content of dry biomass in a liter of medium);

*X*_*0*_ - the initial value of one of the growth criteria.

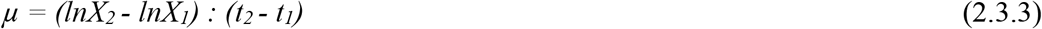

where *X*_*2*_ is the content of dry biomass in a liter of medium at time *t*_*2*_;

*X*_*1*_ – the content of dry biomass in a liter of medium at time *t*_*1*_.

At the final stage, examinations were made to study the growth characteristics of root crops in vitro. To characterize the growth of root crops in vitro, the growth index was used. Statistical data processing was performed using the Microsoft ® Excel computer program. The text shows the arithmetic averages of the parameters. The bars in the diagram correspond to the maximum values of confidence intervals at a 95 % probability level according to the Student’s t-criterion. All experiments were performed at least three times.

## Results and discussion

For the obtained selected steadily growing callus cell cultures, a preliminary analysis of the growth of the cultures was performed and the growth index was calculated. The results of the study of the growth index are presented in Table 1.

**Table 1.** Results of the study of the growth index of the obtained callus cultures

| Callus cell culture | Duration of the subcultivation cycle, day | Dry biomass growth index |
| --- | --- | --- |
| *Scutellaria baicalensis* | 32–37 | 9,4–11,7 |

The main growth characteristics of suspension cultures of medicinal plants are calculated, they are presented in Table 2.

**Table 2.** Results of studying the growth characteristics of suspension cultures cultivated in flasks

| Name of the cell culture | $M_{\max\_dw}$, г/л | $v$, % | $\mu_{dw}$, $cyT^{-1}$ | $I_{dw}$ |
| --- | --- | --- | --- | --- |
| *Scutellaria baicalensis* | 6,16–7,51 | 66,00–85,00 | 0,10–0,12 | 5,87–6,68 |

\* $M_{\max\_dw}$ – maximum value of cell biomass concentration by dry weight; $v$ – cell viability; $\mu$ -specific growth rate; $I$ -growth index.

The complex analysis of the obtained results made it possible to identify the characteristic features of the studied cell cultures.

All lines are represented by two types of cells – mostly meristemoid and some parenchymal-like. Moreover, the number of the latter increased by the end of the stationary phase of the subcultivation cycle in all variants.

For suspension cultures of cells of the medicinal plant Baikal skullcap (Scutellaria baicalensis), the presence of a significant number of cells of an elongated curved shape was noted (Figure 1).

**Figure 1.**
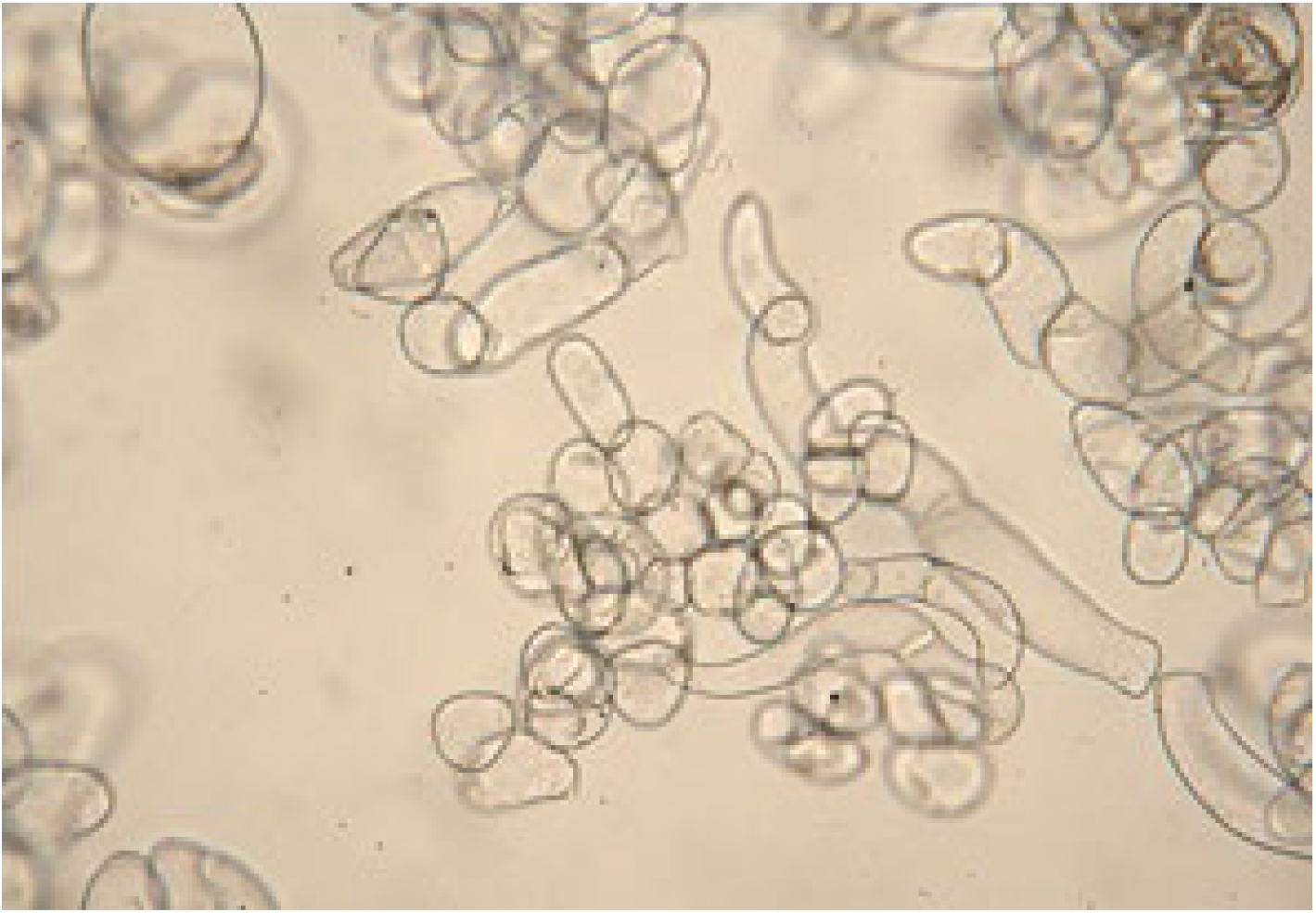
Suspension culture of cells of the medicinal plant Baikal skullcap (Scutellaria baicalensis) under a microscope

The shape of the aggregates is mostly rounded for all lines. According to the degree of aggregation of cells in the suspension, the line of the Baikal skullcap (*Scutellaria baicalensis*) is highly and very coarsely aggregated – more than 60% of the suspension is accounted for by aggregates >0.5 cm in size, while the number of viable single cells and small aggregates (up to 5 cells) was insignificant.

For the studied cell cultures with a control initial planting density of about 0.9–1.5 g/l for dry cell biomass, the subcultivation cycle for determining growth characteristics was 20-22 days. By the 21st day of cultivation, some variants were already in the stage of degradation, while others were still in the stationary phase of growth. During cultivation, all suspensions are characterized by an increase in cell size and the ratio of raw weight to dry weight by the end of the subcultivation cycle, which is associated with an increase in vacuoles during cell aging.

It should be noted that in general, primary suspensions are characterized by extreme instability of growth, significant heterogeneity of the cell population in shape and size. For all the obtained lines of suspension cell cultures, it was noted that a decrease in the volume of inoculum leads to an increase in the duration of the time period for reaching the maximum value by weight of the biomass. In some cases, a decrease in the initial density from 1.0-2.0 to 0.5–0.7 g / l also led to a significant decrease in growth indicators and viability, up to a stop in growth. For this reason, work continues to further optimize the growing conditions.

At the final stage of the task of studying the growth characteristics, root cultures of *Scutellaria baicalensis* were studied *in vitro*.

To characterize the growth of root crops in vitro, the growth index was used. The results of the research are shown in Figure 2.

**Figure 2.**
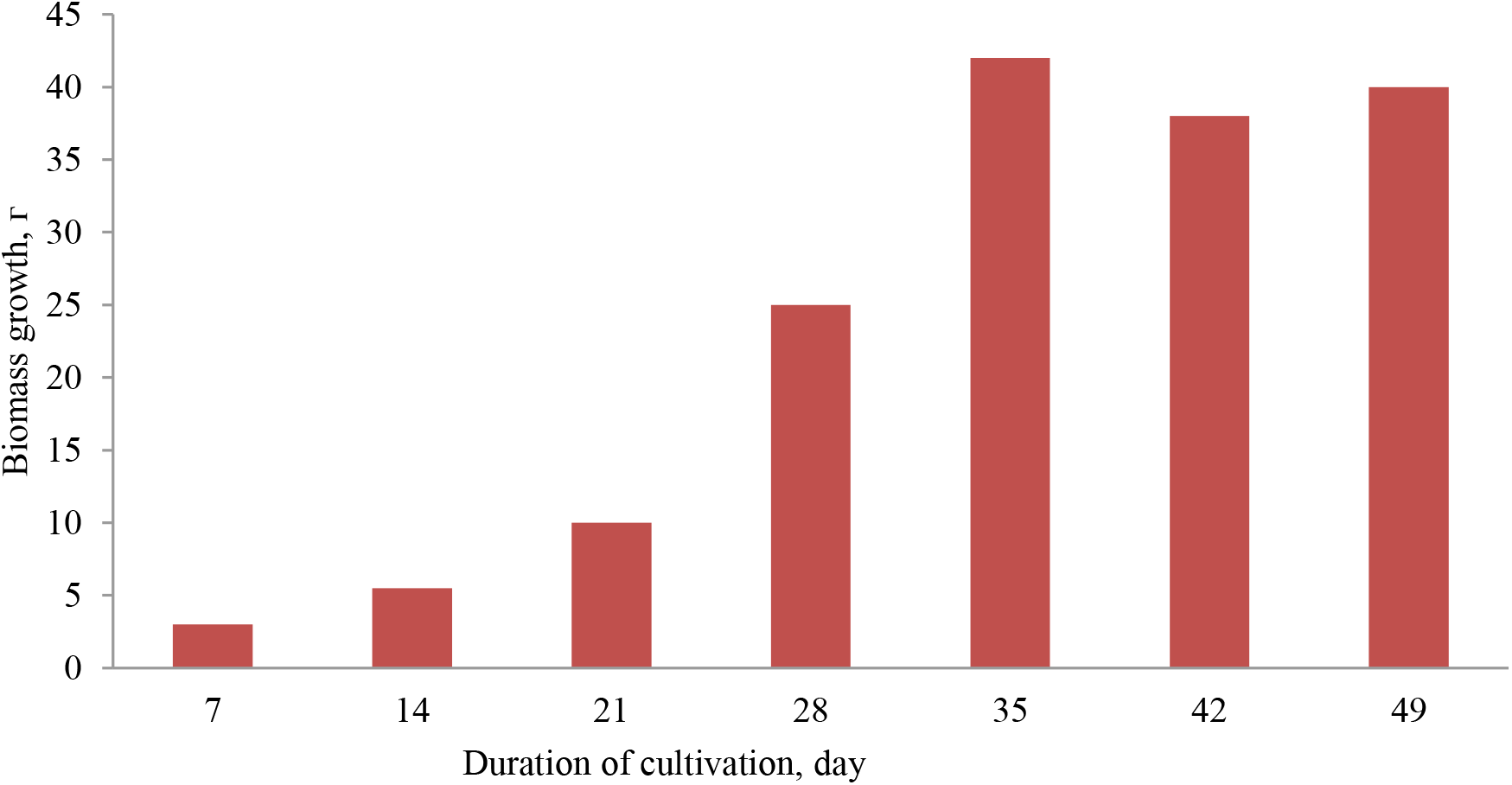
Dependence of the growth index of the root culture in vitro of the Baikal skullcap (*Scutellaria baicalensis*) on the duration of cultivation

The root culture in vitro of the medicinal plant ***Scutellaria baicalensis*** is characterized by intensive growth. The growth index at the end of the cultivation cycle was 40. The growth curve has a standard S-shape, with pronounced growth phases. The stationary phase was observed from 5-7 weeks of cultivation, after which the degradation stage began, at which the rate of biomass growth was not observed (Figure 3.6.3). The root culture growth index was 22.

